# High-density recording reveals sparse clusters (but not columns) for shape and texture encoding in macaque V4

**DOI:** 10.1101/2023.10.15.562424

**Authors:** Tomoyuki Namima, Erin Kempkes, Polina Zamarashkina, Natalia Owen, Anitha Pasupathy

**Author notes:** Corresponding author: Anitha Pasupathy, 1959 Pacific Street NE, HSB G-520, University of Washington Mailbox 357420, Seattle, WA 98195. **Declaration of interests** The authors declare that they have no competing interests. **Author contributions** A.P. and T.N. conceptualized the study. T.N., P.Z., E.K, and N.O. performed the experiments. T.N. and A.P. analyzed the data and wrote the manuscript. P. Z. date of death: July 2020.

## Abstract

Macaque area V4 includes neurons that exhibit exquisite selectivity for visual form and surface texture, but their functional organization across laminae is unknown. We used high-density Neuropixels probes in two awake monkeys to characterize shape and texture tuning of dozens of neurons simultaneously across layers. We found sporadic clusters of neurons that exhibit similar tuning for shape and texture: ∼20% exhibited similar tuning with their neighbors. Importantly, these clusters were confined to a few layers, seldom ‘columnar’ in structure. This was the case even when neurons were strongly driven, and exhibited robust contrast invariance for shape and texture tuning. We conclude that functional organization in area V4 is not columnar for shape and texture stimulus features and in general organization maybe at a coarse scale (e.g. encoding of 2D vs 3D shape) rather than at a fine scale in terms of similarity in tuning for specific features (as in the orientation columns in V1). We speculate that this may be a direct consequence of the great diversity of inputs integrated by V4 neurons to build variegated tuning manifolds in a high-dimensional space.

**Significance Statement:** In primary visual cortex of the macaque monkey, studies have demonstrated columnar functional organization, i.e. shared tuning across layers for stimulus orientation, spatial frequency, ocular dominance, etc. In mid and higher level visual form processing stages, where neurons exhibit high-dimensional tuning, functional organization has been harder to evaluate. Here, leveraging the use of the high-density Neuropixels probes to record simultaneously from dozens of neurons across cortical layers, we demonstrate that functional organization is not columnar for shape and texture tuning in area V4, a midlevel stage critical for form processing. Our results contribute to the debate about the functional significance of cortical columns providing support to the idea that they emerge due to one-to-many representational expansion.

## Introduction

More than six decades ago, Mountcastle and colleagues first discovered cortical “columns” in the somatosensory cortex of the cat, where neurons responsive to stimulation of different peripheral receptors were organized in bands spanning all cortical layers (Mountcastle, 1957). Since that groundbreaking discovery, scientists have demonstrated columnar structure in many cortical regions in a variety of species. Here we use “columns” in a broad sense, focusing on evaluating whether organization runs across layers without any claim of discreteness. In the primate visual system, columns for ocular dominance, stimulus orientation and spatial frequency have been demonstrated in primary visual area (V1) (Albright et al., 1984; DeAngelis and Newsome, 1999), and for motion direction (Albright et al., 1984) and disparity (DeAngelis and Newsome, 1999) in the middle temporal visual area (MT). These demonstrations of columnar structure as well as its absence in some species (Jang et al., 2020) have stimulated intense debate about anatomical origin and functional significance (Horton and Adams, 2005; Purves et al., 1992): do columns represent a critical functional unit or do they result simply from representational expansion (Jang et al., 2020; Ringach 2004, 2007), i.e. a small number of inputs projecting onto a large number of neurons? Here we probe area V4, a midlevel stage in the primate ventral stream, both to gain insight into its functional architecture and to contribute to the debate on the significance of columns.

Columnar structure has been hard to evaluate in the mid- and higher cortical stages of the ventral visual stream critical for form processing. This is primarily because of technical challenges. Unlike tuning for motion direction or orientation, detailed characterization of shape selectivity often requires 100s of stimuli. This makes sequential study of neurons along a recording track, needed to assess columnar structure, difficult because of constraints on experimental time in the awake animal. In the face of this hurdle, studies in V4 and inferior temporal cortex have typically used optical imaging, and more recently multiphoton imaging, to evaluate clustered tuning at a categorical level, e.g. preferred tuning for curved vs rectilinear stimuli or 2D vs 3D shapes, and across the cortical sheet rather than across laminae (Hu et al., 2020; Jiang et al., 2021; Li et al., 2013; Lu et al., 2018; Sato et al., 2009; Srinath et al., 2021; Tang et al., 2020; Tanigawa et al., 2010). Here, for the first time, we leverage the use of high-density Neuropixels probes to examine the fine-scale columnar structure, namely the similarity in the profile of the tuning curve, for shape and texture encoding in cortical area V4 of the awake primate.

Many neurons in area V4 encode information about the shape and texture features of visual stimuli (Desimone and Schein, 1987; Gallant et al., 1993; Kim et al.,2019; Kobatake and Tanaka, 1994; Okazawa et al., 2015; Pasupathy and Connor, 1999). To determine if neurons with similar shape and texture selectivities are clustered across cortical laminae, we probed neuronal responses with a set of 2D shape silhouettes and textures. We assessed similarity in tuning for shape/texture by quantifying the correlation in responses of all pairs of simultaneously studied neurons. To evaluate whether a lack of similarity in tuning was due to tuning dependence on stimulus contrast or stimulus position, we presented our stimuli at two luminance contrasts relative to the background and at multiple positions within the aggregate receptive field (RF). Our results reveal that similarity of shape and texture tuning across laminae is rare in V4: even neurons that exhibit strong shape/texture selectivity that is invariant across contrasts are surrounded by neurons with different tuning preferences.

## Materials and Methods

### Animal preparation and recording sites

Two rhesus macaques (Macaque mulatta) weighing 6.0 kg (female monkey M1) and 9.2 kg (male monkey M2) participated in the experiments. Animals were surgically implanted with custom-built head posts attached to the skull with orthopedic screws. After behavioral training, a metal ring was chronically implanted on the skull surface of each monkey, followed by a craniotomy in a subsequent surgery and the installation of a metal/plastic recording chamber. Since we performed acute Neuropixels probe insertion via native dura each day (see below) it was critical to maintain a thin dural layer with periodic (once every 2-3 weeks) dural debridement under Ketamine (with pain medication). All animal procedures conformed to National Institutes of Health guidelines and were approved by the Institutional Animal Care and Use Committee at the University of Washington.

Positioning of the recording chamber was guided by stereotaxic coordinates based on structural magnetic resonance images taken prior to implant placement, and allowed access to dorsal V4 gyrus between lunate and superior temporal sulci on the right hemisphere. In both animals, locations of probe insertions were visually confirmed post mortem on the cortical surface of dorsal V4 gyrus.

### Experimental design

#### Experimental apparatus and fixation task

During each experimental session, animals were seated in front of a visual display–a liquid crystal display monitor (24 inches; 100 Hz frame rate; 1920 × 1080 pixel size, XL2430-B, BENQ) calibrated with spectrophotoradiometer (PR650; PhotoResearch)--at a distance of 55 cm (M1) or 52 cm (M2). Visual stimulus presentation and behavioral tasks were controlled by custom software written in Python (Pype2, Mazer 2013). Eye position was monitored with a 1 kHz infrared eye-tracker (Eyelink 1000, SR Research). Stimulus onset times were based on photodiode detection of synchronized pulses at the lower left corner of the display monitor (25 kHz sampling rates).

During data collection in this study, animals were engaged in a passive fixation task. Each trial began with the presentation of a white fixation dot (FP; 0.1 degree). Animals were required to maintain eye position within a fixation window of 1.1 degrees for a total duration of 1.6-3.2 sec for water or juice reward. As animals maintained fixation, a sequence of 4-8 stimuli were presented, each for 200 ms, separated by 200 ms inter-stimulus intervals. The first/last stimulus was preceded/followed by a 100 ms blank interval.

### Electrophysiology

#### Probe and Data acquisition configuration

Neuropixels 1.0 and Neuropixels NP1010 (IMEC) were used for recordings in M1 and M2, respectively. Neural signals from the probe and non-neural event signals from other devices (eye-tracker, sync pulse generator, and photodiode) were acquired using PXIe acquisition module (PXIe_1000, IMEC) and multifunction I/O module (PXI-6224, National Instruments-NI), and transmitted to data acquisition Windows computer via the PXI Chassis (PXIe-1071, NI) (Putzeys et al., 2019). Action potential (AP) and local field (LF) signals from the 384 probe contacts (AP: 30 kHz sampling rates; LF: 2.5 kHz sampling rates) were amplified and bandpass-filtered (AP: 0.3-10 kHz; LF: 0.5-500 Hz), and stored for offline analysis (spikeGLX, Janelia Research Campus). Square-wave sync pulses generated with a microcontroller board (Arduino R3, Arduino) were stored in both the neural and non-neural event data recording streams so that spike timing and photodiode detection could be synchronized in offline analysis.

#### Probe insertion

We tracked our probe insertion locations using a custom-built plastic grid with 1 mm spacing. Each day, we marked the target insertion location with india ink on the dural surface using a needle inserted through the grid. We then removed the needle and grid, stabilized the dural surface with a custom-built wire presser foot held by an ultracompact micromanipulator (MO-903B, Narishige), and used one of two methods to insert the Neuropixels probe. For recordings in M1 with the rodent probe (Neuropixels 1.0), which is too fragile to penetrate primate dura, we created a dural eyelet by inserting a short guide tube (3-4 mm length of 27G hypodermic needle, BD) 1-2 mm into the dura and then carefully threading the probe through the eyelet under magnification; the probe itself was mounted on a holder (uMp-NPH, Sensapex) which was mounted on a hydraulic microdrive (MO-97A, Narishige). The short guide tubes were custom made with a blob of epoxy on one end to serve as an anchor against the dura. We used a custom-built guide tube holder and the hydraulic microdrive (MO-97A, Narishige) to insert the short guide tube into the dura and retract the holder to create the dural eyelet. For recordings with the primate probe (Neuropixels NP1010) in M2, we first sharpened the probe using a micropipette grinder (EG-45, Narishige) and then inserted the probe through native dura. In both methods, the experimenter monitored signals from the probe on spikeGLX activity map as the probe was being inserted. Probe reference and ground were connected to the head post of the animal prior to probe insertion. See Namima et al., (2023) for more detailed insertion procedure.

### Preliminary characterization

At the start of each recording session, we used a hand mapping procedure to identify the aggregate receptive field (RF) of neurons recorded along the length of the probe (Figure 1A). Using a gray rectangular bar stimulus of variable length, orientation, aspect ratio, and luminance contrast relative to the background, we mapped the outer boundary of evoked responses from a single recording channel, fit a circle to the mapped RF boundary and the center of the fitted circle was considered to be the RF position. We performed this manual RF mapping for 2-4 channels. choosing channels that were spatially well segregated along the length of the probe, and were associated with a large signal amplitude on spikeGLX AP activity map as well as visually recognizable waveforms on AP spike trace view. The average of the mapped RF locations across the sampled channels was referred to as the “aggregate RF center”, and was used as the center location for the presentation of visual stimuli.

**Figure 1.**
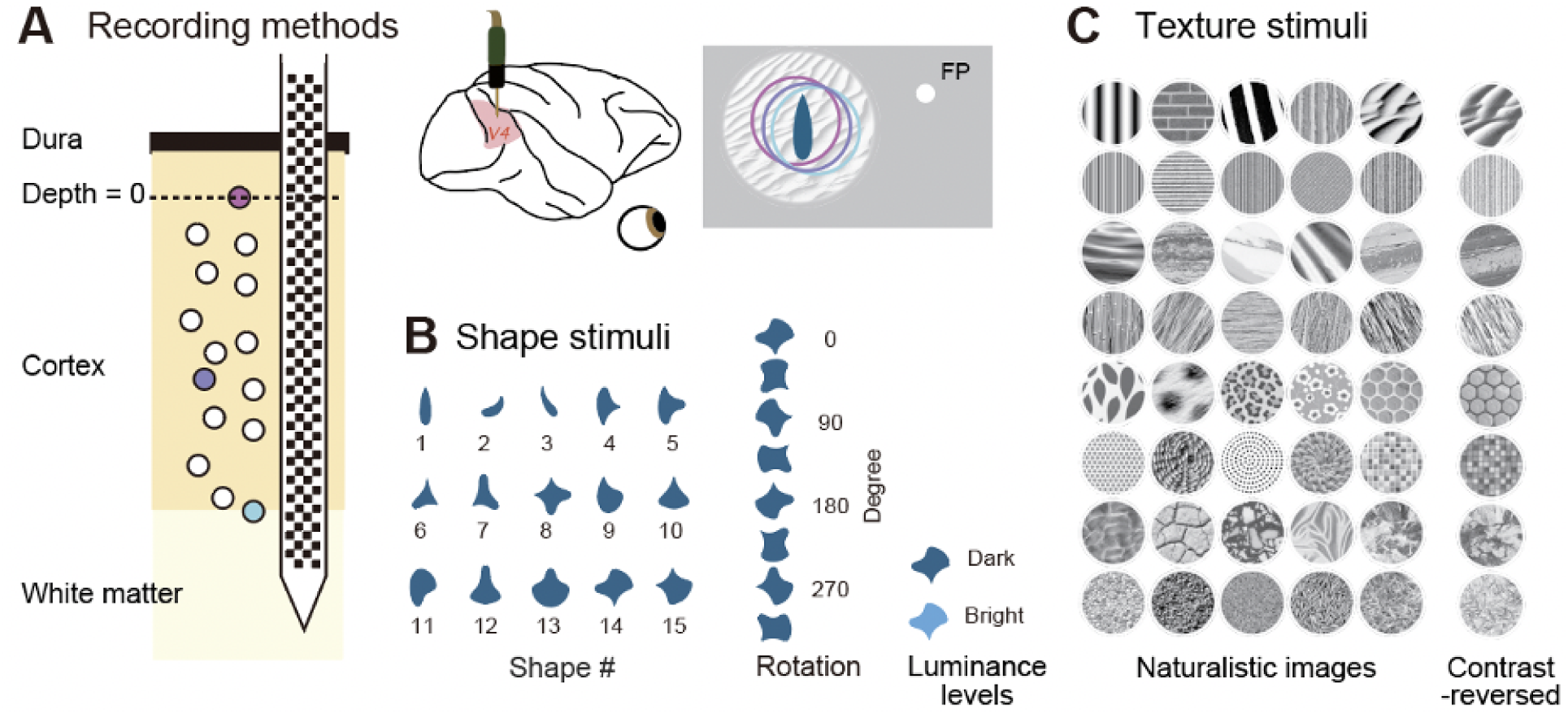
Recording methods and visual stimuli. **(A)** A high-density Neuropixels probe (left) was used to study the responses of subpopulations of V4 neurons across cortical laminae. We targeted dorsal V4 on the prelunate gyrus (middle panel). Neurons across laminae (dots in the left panel) had partially overlapping RFs (circles in the right panel) identified during preliminary RF characterization. Visual stimuli were presented at the center of the aggregate RF. FP: fixation point. **(B)** Visual stimuli included 15 two-dimensional shape silhouettes, each presented at 8 different rotations in 45° increments. Shapes were presented at two luminance contrast levels (bright or dark) relative to background. **(C)** Forty naturalistic textures at different orientations on each trial (see Methods) were presented in two versions: original or contrast-reversed (compare last two columns).

### RF localization

To facilitate more precise post-hoc characterization of the RF location of individual neurons along the length of the probe, we conducted an automated RF localization experiment. As animals fixated, visual stimuli were presented on a 31 × 31 or 41 × 41 square grid, centered on and scaled to cover the aggregate RF identified during preliminary characterization. Each grid location was sampled once, with either a shape or a texture stimulus chosen randomly from our main stimulus sets described next. The size was common across the grid locations in the same penetration. Stimuli were presented for 80 or 100 ms duration with a 100 ms interstimulus interval.

### Visual stimuli

During the main experiment, visual shape and texture stimuli were presented against an achromatic background (8 cd/m^2^; x = 0.33 and y = 0.33 in CIE-xy chromaticity coordinates) at the aggregate RF center (see Preliminary characterization). The shape set included 15 simple geometric shapes each presented at 8 different rotations in 45° increments, resulting in a total of 120 shape stimuli (Figure 1B; also see Pasupathy and Connor, 2001). For this study we presented shapes filled with a uniform cyan (x = 0.196, y = 0.188), either darker or brighter than the background luminance (4 cd/m^2^ or 12 cd/m^2^). The shapes were typically sized to fit entirely within the estimated V4 RF scaled for eccentricity (based on Gattass et al., 1988; see Pasupathy and Connor, 1999) except for five experiments in M1 where the shapes were larger in order to overlap a greater fraction of the RFs of the neurons being recorded.

The texture stimulus set was composed of 40 naturalistic grayscale textures (Figure 1C) with pixel values ranging from 1 cd/m^2^ to 30 cd/m^2^ (mean luminance 13 cd/m^2^). Textures were presented either in their original version or as contrast-reversed images, where pixel luminance contrast was inverted relative to the mean luminance of 13 cd/m^2^. All texture images were presented through a circular aperture with a blurred boundary and were sized to cover the area occupied by the RFs of multiple single neurons characterized each day during manual mapping (see Preliminary characterization). On each stimulus presentation, texture images were presented at one of 8 randomly chosen rotations at 45° increments to assess similarity in tuning for higher order texture statistics rather than tuning for local oriented features. For both experiments, we included blank stimulus periods to assess baseline activity.

To evaluate whether scatter in the RF position across neurons contributed to the sparse clustering observed in the shape experiments, we conducted a position control test. We used the same 15 shapes (Figure 1B) each at a single orientation, with the shape orientations chosen to provide roughly even sampling of curvatures across angular position. Each stimulus was presented in 5 positions: at the aggregate RF center and at four positions displaced ½ × RF diameter (north, south, east, and west) relative to the aggregate RF center. In the position control experiment, shape stimuli were evenly filled with dark cyan (4 cd/m^2^) and were sized as in the main experiment.

### Dataset and data inclusion criteria

We conducted 30 and 44 sessions of main shape test in M1 and M2, respectively. The position control experiment with shape stimuli was conducted in 33 of 44 sessions in M2 only. The texture test was conducted in 17 and 34 sessions in M1 and M2, respectively. Shape and texture tests were conducted in different experimental sessions in M1. In M2, we conducted shape and texture tests in 34 sessions, of which 30 sessions included 6 or more neurons that were visually driven by both shape and texture stimuli; these sessions were used to compare invariance in shape and texture tuning across stimulus contrasts. The automated RF localization experiment was conducted in all recording sessions. Even when all four tests were conducted in the same penetration, they were conducted in separate blocks. Across experiments, the median number of repetitions for individual shape stimuli was 5 (minimum = 3) and for texture stimuli and position control was 8 (minimum = 4). All clusters deemed “good” after automatic and manual curation (see Data preprocessing below) were included in our analysis to identify the most superficial neuron (see Depth from superficial neuron). For assessments of tuning similarity, we excluded neurons that were not visually driven. For this we used a very liberal definition of responsiveness: each neuron had to exhibit a response that was ≥ 5spikes/sec above baseline for one shape or texture stimulus at either luminance contrast level. This excluded 461 and 366 neurons across our dataset for shape and texture analyses, and left 1850 and 979 neurons respectively. Thus, a very liberal criterion excluded roughly 20% of data from our shape test and 27% from our texture test. These excluded neurons presumably require other stimulus dimensions (motion, 3D cues, etc.) and/or other shapes and textures not tested here.

### Data preprocessing

The flow chart in Figure S9 summarizes the preprocessing steps. Each day, binary data files collected across experiments were combined into a single binary file using custom-built MATLAB code (Mathworks Inc.). The combined binary files were then processed with open source automated spike sorting software (high pass filtering with 300 Hz; Kilosort 2.0 (Pachitariu et al., 2023), or Kilosort 2.5 (Pachitariu et al., 2023; Steinmetz et al., 2021)) with default parameters, followed by manual curation with open source software (phy2 template-gui, cortex-lab). During manual curation, we categorized Kilosort-detected waveform clusters as “good”, “multi-unit activity (mua)” or “noise”. Waveform clusters associated with low firing rate (< 0.1 spikes/sec) and/or low amplitude waveforms (almost flat waveforms) were classified as “noise”. Clusters with waveforms that were large in amplitude, with minimal high frequency fluctuations across time, and distinct in shape from other clusters isolated on nearby electrode channels, were categorized as good clusters (also see Bigelow et al., 2023). Because Kilosort allows overlapped spikes to be fit with different templates, some good clusters may be double-counted (because the residuals may be fit with a second template). We identified this duplication based on the similarity of the inter spike interval (ISI) histogram and the cross-correlogram and excluded one of the two clusters from our dataset. Specifically, we examined the cross-correlogram between pairs of putative good clusters and excluded one cluster as a double-count if the cross-correlogram exhibited a zero-centered distribution with a very large, narrow peak. All other waveform clusters were categorized as “mua”. Finally, we examined the ISI distribution of good clusters and rejected a subset when > 5% of the ISIs violated the 2 ms refractory period and/or the mode of the ISI histogram was less than 2 ms. All remaining waveform clusters labeled as “good” were included in all further analyses.

After automatic and manual sorting, one final pre-processing step involved the alignment of the different data recording streams. Because the neural and non-neural event signals were stored in separate streams with independent clocks with rates that could vary with temperature, we linearly regressed the sync pulse edge times stored in the two streams to bring the spike times, event timestamps and photodiode signals into register. Analyses of spike times were performed using custom-built MATLAB code using freely-distributed MATLAB toolboxes (Spikes, cortex-lab; npy-matlab, Kwik Team).

### Data analysis

#### Firing rate computation

Spike counts from single neurons in response to individual stimuli were computed within a 30-200 ms window after stimulus onset. Then, firing rates (FRs) averaged across stimulus repetitions were computed. Baseline FRs were similarly computed during blank stimulus presentations.

#### Depth from superficial neuron

For both probes used in these studies, the default channel maps of recording contacts are organized into three banks, and during each recording session data may be collected from one bank of 384 contacts. We used the deepest bank (Bank0, closest to the tip of the probe) for all recording sessions. To estimate the depth of individual neurons from the cortical surface, we first assessed the axial position of each neuron along the probe by computing the center mass of the waveform template across all channel positions where the waveform was detected (templatePositionsAmplitudes.m, Spikes toolbox). The depth of each neuron was then calculated as the relative distance between its position and that of the most superficial neuron along the probe length.

#### Current source and sink profile along the probe

We conducted current source density (CSD) analysis to estimate the position of the cortical input layer along probe length. First, for individual channels, we down-sampled local field (LF) signal from 2.5 kHz to 1 kHz and then subtracted the mean from the down sampled LF signal. The ground-subtracted LF signal was transformed from 16-bit data format (i) to actual voltage unit (V). Then, we applied a 3^rd^ order Butterworth bandpass filter with a range of [0.5 100] Hz to the LF signal. LF signal to first stimulus presented in each trial was averaged across multiple trials and smoothed using a Gaussian function across adjacent 10 contacts. For the trial averaged LF signal, we computed standard current source density (Freeman and Nicholson, 1975): second derivative between LF signals from neighboring 3 contacts. The CSD were calculated separately for four channel columns along the probe length but one of CSDs were visualized. Our CSD visualization was accomplished by applying a Gaussian smoothing with surrounding 10 contacts and a sign-inversion over the calculated CSD values.

#### Tuning similarity and contrast invariance

We computed the correlation coefficient between responses of two neurons to assess similarity in tuning, and between responses of a neuron to two stimulus types (e.g. bright and dark) to assess contrast invariance in tuning across stimulus attributes. However, when the number of stimulus repeats are low, Pearson’s correlation coefficient can be heavily biased, and this bias is related to the trial-to-trial variability of neuronal responses (see detail in Pospisil and Bair 2021). We corrected the attenuation in correlation using methods developed by Pospisil and Bair. Briefly, we measured spike counts on individual stimulus repeats and used a square root transform to stabilize variance. We then computed the numerator and denominator of the correlation coefficient squared, estimated, and subtracted the bias terms from the numerator and the denominator separately, divided the bias-corrected numerator by the denominator and took the square root of the quotient to calculate the noise-corrected correlation coefficient. Finally, the sign of the noise-corrected correlation coefficient was set to that based on the Pearson’s correlation coefficient and the values were truncated to lie in the [-1 1] range. We assessed a confidence interval for each noise-corrected coefficient by conducting bootstrap resampling (number of bootstraps = 500) with repetition of spike counts on individual stimulus repeats. The noise-corrected coefficient was considered unreliable if the confidence limit was > 0.5, which is the case when the signal-to-noise ratio is weak.

#### Clusters of similarly tuned neurons

To assess tuning similarity among neighboring neurons, we related tuning similarity between a neuron and its neighbors to interneuron distance by linear regression (fit.m, MATLAB). We assessed the significance of the fit at the 95% confidence level (confint.m, MATLAB). If tuning similarity is high for nearby neurons, and similarity declines with distance, we would expect a positive y-intercept (*I_similarity_*) and a negative slope (*S_similarity_*). Nearby neurons with dissimilar tuning would be associated with an intercept close to zero. We excluded some neurons from our clustering analysis (shape: n = 1/1850, texture: n = 74/979) because we had too few data points (two or less) of tuning similarity at a given contrast level and thus we were unable to quantify confidence limits for slope and intercept of linear regression fit. This exclusion was necessary because for assessing tuning similarity, we included all neurons that responded to either bright or dark stimuli (see Dataset and data inclusion), but tuning similarity was evaluated separately for bright and dark stimuli. Therefore, the tuning similarity estimates for one contrast could be unreliable when both neurons of the pair were not driven or exhibited poor signal-to-noise ratio (see white pixels in Figure 3). If this was the case for a neuron when paired with many others within a penetration, either the regression line per se or the confidence limits could be un-computable for a neuron.

### Statistical tests

To assess whether clustering of selectivity based on responses at two luminance contrasts were similar, we computed the Spearman’s rank correlation between the y-intercept of individual neurons at the two luminance contrast levels.

To compare tuning invariance for shape vs texture within individual penetrations where we studied both, for each penetration we computed the mean and standard deviation of tuning invariance of all recorded neurons within a penetration, separately for shape and texture. Then we assessed the linear relationship between the means (and standard deviations) for shape and texture by computing the Pearson’s correlation coefficient. Wilcoxon rank sum test was conducted to evaluate whether there was a statistically significant difference in shape/texture tuning invariance between subpopulations of neurons with and without significant intercept.

### RF localization

To estimate the location of each neuron’s RF, we computed the firing rate of neurons at each grid location during stimulus presentation (spikes/sec; either 0-80 ms or 0-100 ms after stimulus onset), smoothed the firing rate over a 3 × 3 grid (conv2.m, MATLAB) and subtracted the baseline FR. After truncating negative values to zero, the smoothed RF maps were fitted with 2D Gaussian kernels to identify the RF center. For each neuron, the RF fit was considered reliable only when the mean FR over a 3 × 3 grid centered on the peak was > 5 spikes/sec. In the position test, we aimed to determine the optimal stimulus location for each neuron. We calculated the dispersion of responses across 15 shapes at five locations relative to the aggregate RF center (center, north, south, east, and west of aggregate RF) by computing the ratio of variance to mean of the FRs across stimuli at each location. The location with the greatest dispersion for each neuron was referred to as the optimal stimulus location and was used to estimate optimized tuning similarity.

### Simulation

We conducted simulations to visualize the tuning similarity matrices we might observe for different patterns of tuning progression along the length of the probe. We created a sub-population of 30 model V4 neurons that were tuned to boundary curvature as quantified by the angular position × curvature model (Pasupathy and Connor, 2001). Specifically, shape responses of each neuron were dictated by a 2D Gaussian function in the angular position × curvature space. The peak position along the angular position and curvature dimensions were chosen either based on a systematic progression with small fluctuations, or at random (see Results, Figure 10). Peak amplitude and standard deviations were set for individual neurons; standard deviations were set to random values from the normal distribution with μ = 60° (σ = 10) and μ = 0.5 (σ = 0.1) along the angular position and curvature dimensions, respectively. For each of the 30 neurons, we first predicted responses to the 120 shapes in accordance with the model. Then, to simulate noisy neuronal responses, for each shape we constructed 200 ms long spike trains using a Poisson process with mean specified by the predicted response. We computed similarity matrices between all neuronal pairs as described above. Because our stimulated responses were with trial-to-trial variability, we used noise-corrected correlation coefficient to assess shape tuning similarity between model neurons.

## Results

We studied the responses of subpopulations of V4 neurons across cortical laminae using high-density multi-contact probes (Neuropixels, Figure 1A) while monkeys performed a passive fixation task. Each day, we identified an aggregate receptive field (RF) for all neurons simultaneously studied in a penetration (see Materials and Methods), and presented either two-dimensional shape silhouettes (Figure 1B) or naturalistic texture stimuli (Figure 1C) centered within the aggregate RF. From data collected during each recording session, we assessed the similarity in shape/texture tuning between all pairs of recorded neurons. Because V4 neurons are sensitive to luminance contrasts (Bushnell et al., 2011; Desimone and Schein, 1987) we presented stimuli at two luminance contrasts. This allowed us to assess the invariance in shape/texture tuning across contrast reversals for each neuron. Using these metrics (tuning similarity and tuning invariance, respectively) we evaluated whether there were clusters of neurons that exhibit similar shape and texture tuning. We also examined whether any lack of tuning similarity could be attributed to heterogeneity in RF profiles across simultaneously recorded neurons.

### Example penetration: Similarity in shape tuning

Figures 2 and 3 summarize results from one recording session where we observed a cluster of neurons that were similarly shape-tuned. We recorded from 24 well-isolated neurons in this session (Figure 2A, left: #1-#24; superficial to deep) over a ∼2400 μm length of the probe. Current source density analysis (CSD, see Materials and Methods) reveals a current-sink at ∼53 msec after stimulus onset at a recording depth of ∼1073 μm relative to the most superficially recorded neuron in this session. We also confirmed that these simultaneously studied neurons were indeed distinct by comparing their waveforms (Figure 2A, right), response latency (Figure 2C), response strength (Figure 2D) and ISI histograms (see Figure 2A, bottom).

**Figure 2.**
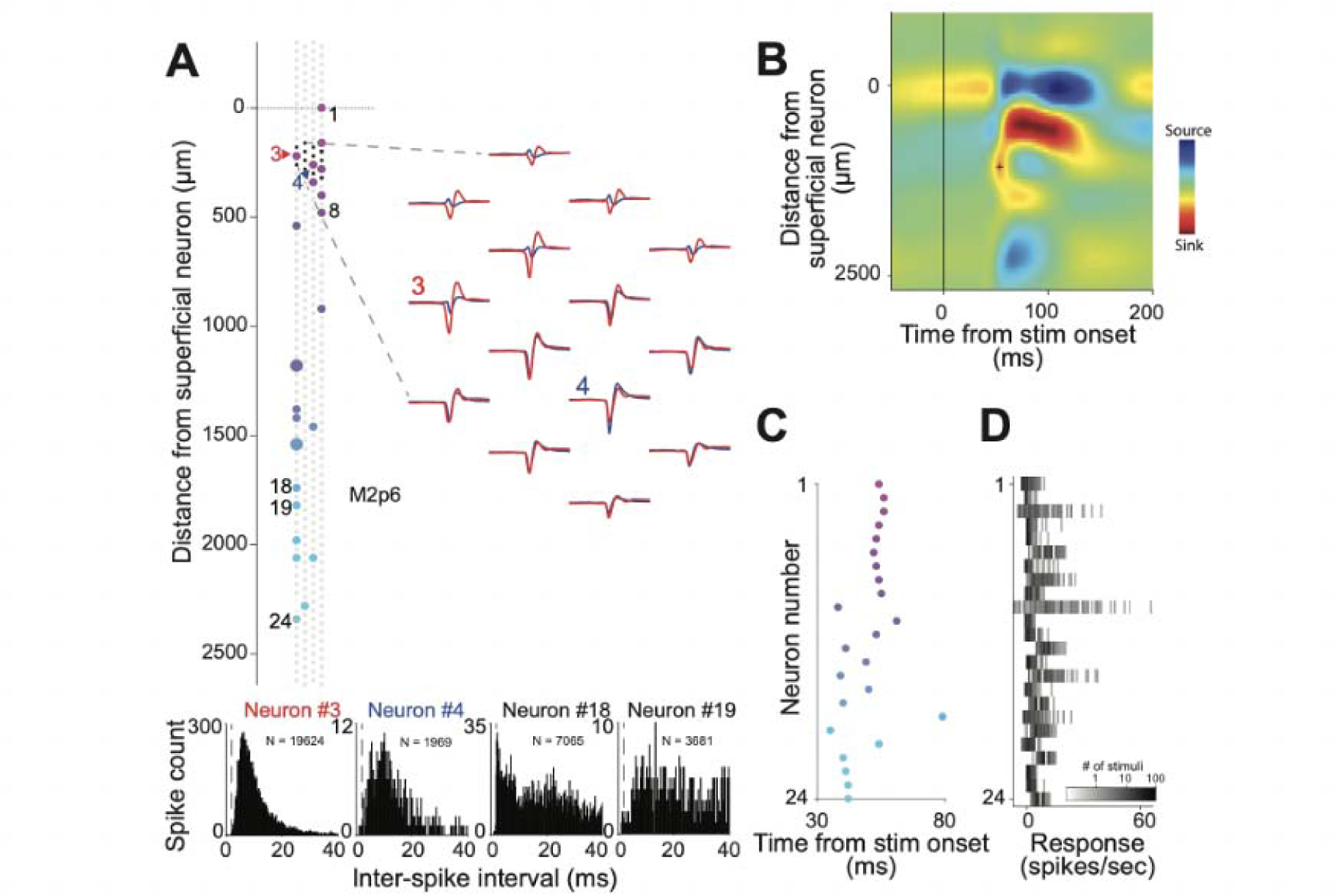
Example recording session: neuronal metrics and CSD. **(A)** Location, waveforms and inter-spike interval (ISI) histograms of recorded neurons in penetration M2p6. *Left.* Location of neurons recorded during this session (n = 24) indicated by the contact closest to the center of mass waveform amplitudes across contacts. Neuron number (color) is rank-ordered relative most superficial neuron. Multiple neurons recorded on the same contact are denoted by larger dots. *Right.* Spike waveforms from two example neurons (#3: red; #4: blue) recorded on the same set of 14 contacts (left: black squares). *Bottom*. ISI histograms (bottom, N: # of spikes; bin size = 0.2 ms) for 4 example neurons based on spikes across the entire recording session. Refractory period (2 ms) is indicated by a dashed line. **(B)** Current source density profile along the probe length. Current-sink (red) and -source (blue) were evaluated by applying standard current source density (CSD) method to trial-averaged local field (LF) signal evoked by shape stimulus onset (see Materials and Methods). Cross marker indicates the current-sink considered to be layer 4 (depth = 1073 μm, time to current-sink = 53 msec after stimulus onset). **(C)** Onset latency of responses to dark stimuli. Latency was quantified as the time to half-peak of the peri-stimulus time histogram (PSTH) constructed from responses to shape stimuli. For each neuron, spike rasters for all dark stimuli were accumulated, and then were smoothed using a Gaussian smoothing window (size of kernel = 4, sigma = 10). Time to half-peak was assessed starting 20 ms after stimulus onset since latencies < 20 ms are likely noise. PSTHs were constructed relative to baseline activity, which was quantified as mean activity during the 30 ms time period before stimulus onset. **(D)** Frequency histogram for the average firing rate (baseline-subtracted, X axis) of each neuron (Y axis) across 120 dark stimuli. Grayscale shows the number of stimuli (log-scale) associated with each firing rate.

**Figure 3.**
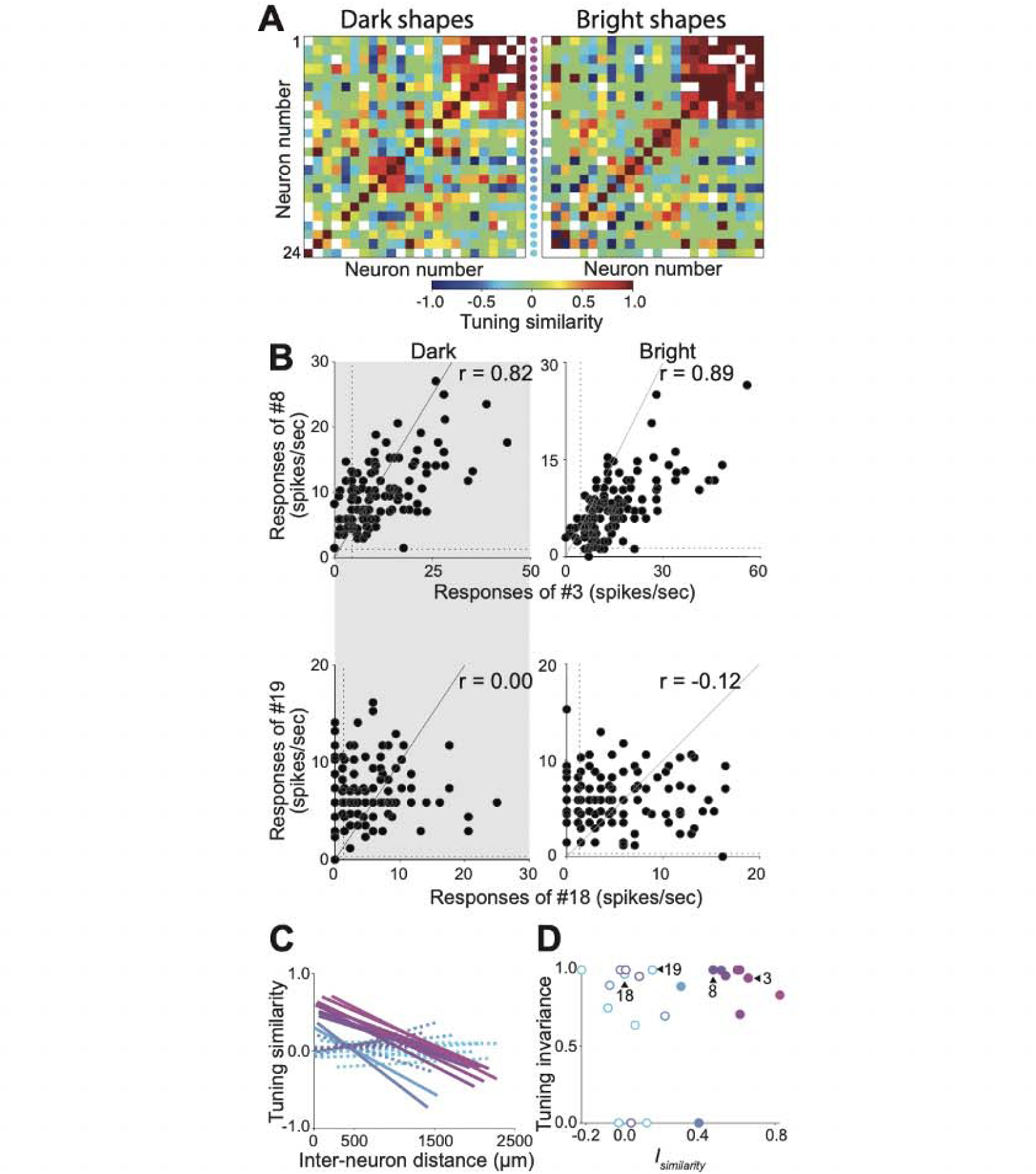
Example recording session: cluster of similarly shape-tuned neurons. **(A)** Tuning similarity in the responses to dark (left) or bright (right) shape stimuli across the set of 24 simultaneously studied neurons in penetration M2p6 (see Figure 2). Similarity values range from -1 (blue) to 1 (red). White pixels denote neuron pairs for which tuning similarity could not be quantified reliably (due to poor signal-to-noise ratio, see Materials and Methods). Neuron number runs from most superficial (1: purple) to deepest (24: cyan) along the probe. **(B)** Scatter plots relating the responses of two pairs of neurons (top: #3 vs #8; bottom: #18 vs #19; see Figure 2A) for dark (left) and bright (right) shape stimuli. The top pair shows high tuning similarity (r = 0.82, left; r = 0.89, right) but the bottom does not (r = 0.00, left and r = -0.12, right) even though all four neurons exhibit a broad range of responses that show high tuning invariance (see Figure 3D). Dotted lines indicate baseline FRs. **(C)** Linear regression lines characterize the trend of tuning similarity (Y axis) versus inter-neuron distance (X axis) for each neuron (line color) and its neighbors (see Materials and Methods). Attenuation of tuning similarity with increasing inter-neuron distance was captured by a positive intercept and a negative slope. Solid and dashed lines indicate regression lines with and without significant intercept. **(D)** Shape tuning invariance (Y axis) versus y-intercept of regression lines in 3C (X axis: *I_similarity_*). Filled and open circles indicate neurons with and without significant intercept, respectively.

Figure 3 illustrates similarity in tuning for all pairs of recorded neurons in this session. We quantified the strength of similarity in terms of the noise-corrected correlation coefficient (Pospisil and Bair 2021, see Materials and Methods) between responses of all pairs of neurons to shapes darker (Figure 3A, left) or brighter (right) than the background. The strength of similarity ranges from -1 (blue: opposite preference) to 1 (red: identical preference) with white denoting a lack of reliable responses (confidence limit > 0.5; see Materials and Methods). These matrices are symmetric; diagonal elements are at 1.0 (correlation between two identical sets of responses) except when signal-to-noise ratio is weak (white pixels). For this recording session, a small group of neurons (#1-#9) spread over a 540 μm extent of the probe in superficial cortex (see Figure 2A, left) exhibited high similarity in their responses to the tested shapes. This was the case regardless of whether the shapes were darker (Figure 3A, left) or brighter (right) than the background. Thus, these neurons may be considered to constitute a functional shape domain of similarly shape-tuned neurons. However, the majority of neurons in this penetration did not exhibit tuning that was similar to their neighbors.

To rigorously quantify the spatial profile of clustering, i.e. how tuning similarity changes along the rows or columns in Figure 3A as a function of inter-neuron distance, we used linear regression to relate tuning similarity between a neuron and its neighbors to the distance between them. Figure 3C shows the regression lines for the 24 recorded neurons based on the responses to dark stimuli; results with bright stimuli (not shown) were similar. For some neurons (neurons #1-9, #14, and #17), regression lines were characterized by a large positive y-intercept (*I_similarity_*) significantly different from zero (p < 0.05, solid lines) and a negative slope (*S_similarity_*). These were neurons that showed high similarity in tuning with nearby neurons (< 600 μm away) that declined with distance. For other neurons, mostly those deeper in cortex (cyan lines), similarity in tuning was weak regardless of distance between neuron pairs; in this case the regression lines were associated with low *I_similarity_*and shallow negative or positive *S_similarity_* that were not statistically different from zero (dashed lines). In the latter group of neurons, lack of statistically significant *I_similarity_* and *S_similarity_*was because nearby neurons failed to show similar tuning (bottom, Figure 3B) and not because we failed to sample nearby neighbors. For example, neurons #3 and #8, recorded on contacts 260 μm apart, share a great deal of similarity in tuning (*r_similarity_* = 0.82), but neurons #18 and #19 recorded on contacts 80 μm apart do not (*r_similarity_* = 0; Figure 3B).

Previous studies have shown that V4 neurons are sensitive to luminance contrasts (Bushnell et al., 2011; Desimone and Schein, 1987), and so it is possible that the lack of tuning similarity may be due to contrast dependence of shape responses or tuning, i.e, some neurons may exhibit differential responses to shape only at a specific luminance contrast. To consider this possibility, for individual neurons we asked whether the shape tuning curve was similar when measured with bright vs dark shapes. We computed the invariance in shape tuning to bright and dark stimuli in terms of the noise-corrected correlation coefficient and related the contrast invariance of shape tuning (Y axis in Figure 3D) to *I_similarity_* (X axis). A high tuning invariance (close to 1 along Y axis) would imply that the neuron exhibits similar tuning to shape regardless of contrast polarity, i.e. the shape tuning is contrast-invariant. For a majority of neurons recorded on this day (19/24 neurons), contrast invariance of shape tuning was high (> 0.5) regardless of the value of *I_similarity_*. For example, neurons #18 and #19 exhibit high contrast-invariant shape tuning but show weak similarity in shape tuning (left, Figure 3D). Furthermore, these neurons showed a broad dynamic range in their responses (bottom, Figure 3B), implying that the lack of similarity in tuning between neurons was not due to poor stimulus-evoked responses nor due to lack of shape tuning. Overall, the example recording session illustrated here demonstrates evidence for a small cluster of neurons that share similar shape tuning.

Unlike results illustrated in Figure 2 and 3, many other penetrations showed no evidence of clustered tuning for shape. Figure 4 summarizes results from one such penetration. Here we studied the responses of 42 neurons over a 2580 μm distance (Figure 4A). Many of these neurons were recorded on nearby contacts, and yet, we did not see any evidence of clustered tuning similarity for either dark (Figure 4B, left) or bright (right) stimuli: while some pairs of neurons exhibit high similarity of tuning (orange/red pixels), they are sporadic and not localized in a cluster along the length of the probe. Lack of clustered tuning similarity was not due to poor stimulus-evoked responses: many neurons showed a broad dynamic range in their responses to shape stimuli (Figure 4C) and peak responses were often high (exceeding 15 spikes/sec in 28/26 neurons for dark/bright stimuli). When we related inter-neuron distance to tuning similarity (Figure 4D) most neurons were associated with low y-intercepts (*I_similarity_*) and shallow slopes (*S_similarity_*) implying that nearby neurons did not share similar tuning. Furthermore, we observed high contrast invariance in the shape tuning of individual neurons (Y axis, Figure 4F) as exemplified by the scatterplots of three example neurons in Figure 4E: response strength depended on luminance contrast, especially for neurons #18 and #32, but the shape tuning was highly contrast-invariant as captured by the strong positive correlation. We found that current-sink with short latency (∼52msec) emerged only in relatively shallower along the penetration (570 of 2580 μm, Figure 4G) but no other sink in deeper cortex, suggesting that this was unlikely a penetration along the sulcal bank. These results imply that the lack of high similarity in tuning was likely caused by a diversity of preferred shape features across simultaneously studied neurons, rather than weak driving by our stimulus protocol and or penetration down the sulcal bank. Thus, Figure 4 provides evidence for a penetration full of neurons with diverse shape selectivity.

**Figure 4.**
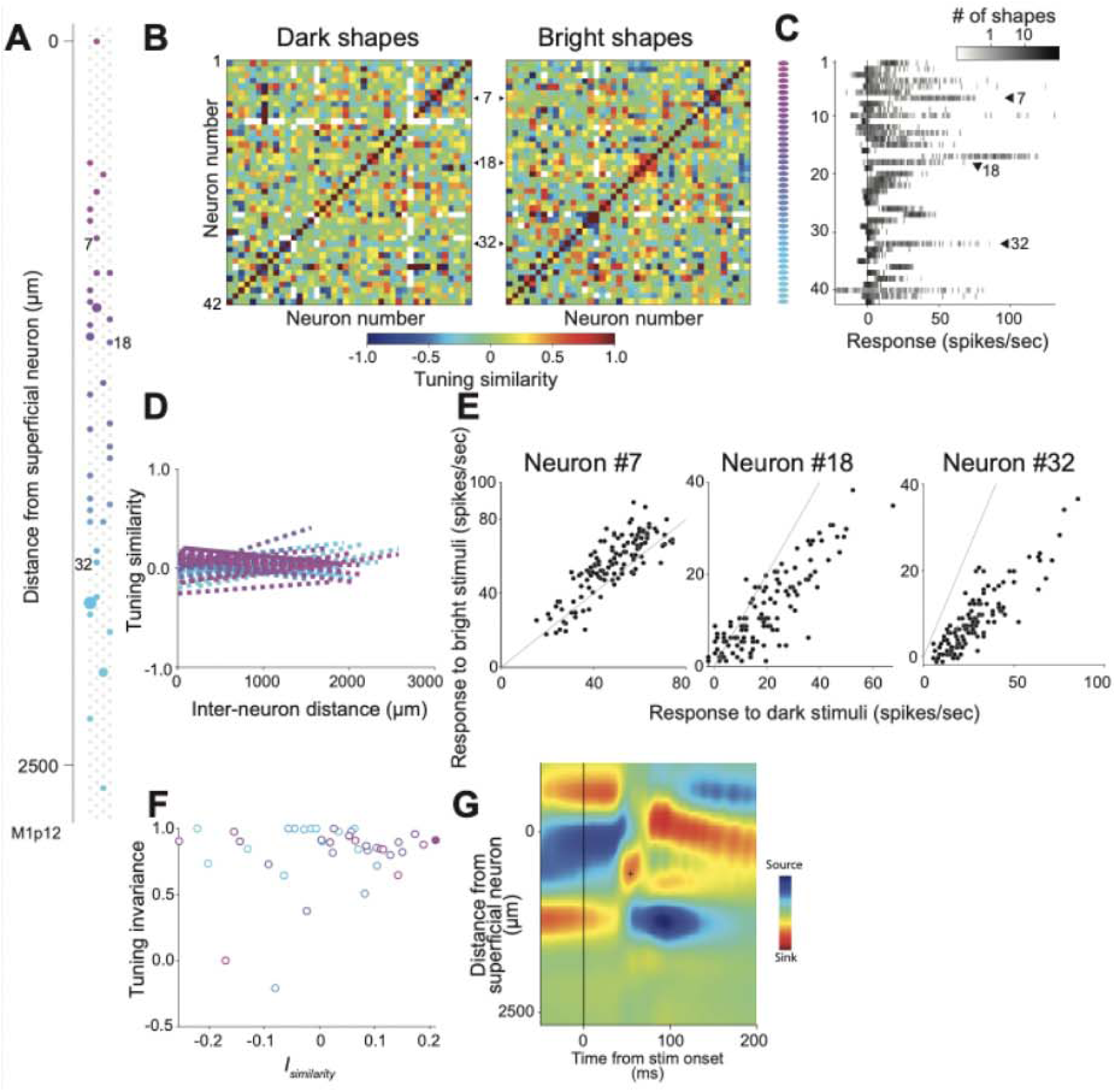
Example session without a cluster of similarly shape-tuned neurons. **(A)** Location of recorded neurons (n = 42) along the probe. Larger dots denote multiple neurons recorded on the same contact. **(B)** Tuning similarity across all pairs of recorded neurons based on the responses to shape stimuli darker (left) or brighter (right) than the background. All other details as in Figure 3A. Evidence of clustered shape tuning is absent. **(C)** Frequency histogram for the average firing rate (baseline-subtracted, X axis) for each neuron (rows) across the 120 dark shape stimuli. Grayscale shows the number of stimuli (log-scale) associated with each firing rate bin. **(D)** Linear regression lines characterizing the trend of tuning similarity (Y axis) versus inter-neuron distance (X axis). All other details as in Figure 3C. Y-intercept for the regression line was significantly different from zero for only one neuron (solid line). **(E)** Responses from 3 example neurons in this session (#7, #18 and #32) for dark (X axis) versus bright (Y axis) shapes. Baseline-subtracted average firing rates are shown. Solid gray lines indicate identity lines. **(F)** Y-intercept of regression lines (X axis: *I_similarity_*) plotted as a function of shape tuning invariance across contrasts (Y axis). All details as in Figure 3D. Neurons in this penetration did not exhibit shape tuning that was similar to their neighbors (filled) but most neurons did exhibit high shape tuning invariance. Only one neuron with Y axis = 0 showed low reliability in the tuning invariance metric. **(G)** CSD profile constructed from LFP signal evoked with shape stimuli. Cross marker on the current-sink (red) considered to be layer 4 was at depth = 570 μm relative to most superficial neuron. Time to current-sink = 52 msec after stimulus onset). All other details as in Figure 2B.

### Population results: Shape tuning

We studied the responses of 1850 neurons across 74 recording sessions (M1: 30; M2: 44), ranging from 6 to 76 neurons per penetration. Figure 5A summarizes how tuning similarity declines with distance across all recorded neurons. As with the examples in Figures 3 and 4, we found a minority of neurons (328/1850; M1:188; M2:140, blue in Figure 5A) with positive *I_similarity_*, significantly different from zero (p < 0.05) and a negative *S_similarity_*, reflecting high similarity with nearby neurons that declined with distance. The majority (n = 1521) had *I_similarity_*that was not significantly greater than zero, reflecting the fact that nearby neurons had tuning that was quite different. Overall, *I_similarity_* and *S_similarity_* were significantly negatively correlated (not shown) and we seldom found neurons associated with positive *S_similarity_*, i.e. similarity that increased with distance: only one neuron (in M2) was associated with a significantly positive *I_similarity_*and *S_similarity_*. We also found that tuning similarity (*I_similarity_*) based on responses to dark and bright stimuli were consistent (Figure 5B). This lack of clustered shape selectivity was despite the fact that contrast-invariant shape tuning was widely prevalent across our dataset: median correlation between responses to dark and bright stimuli in individual penetrations was 0.62 (not shown). Thus, more than half of the recorded neurons were well-driven and exhibited contrast-invariant shape tuning. Thus, lack of high similarity was unlikely due to weak driving across penetrations.

**Figure 5.**
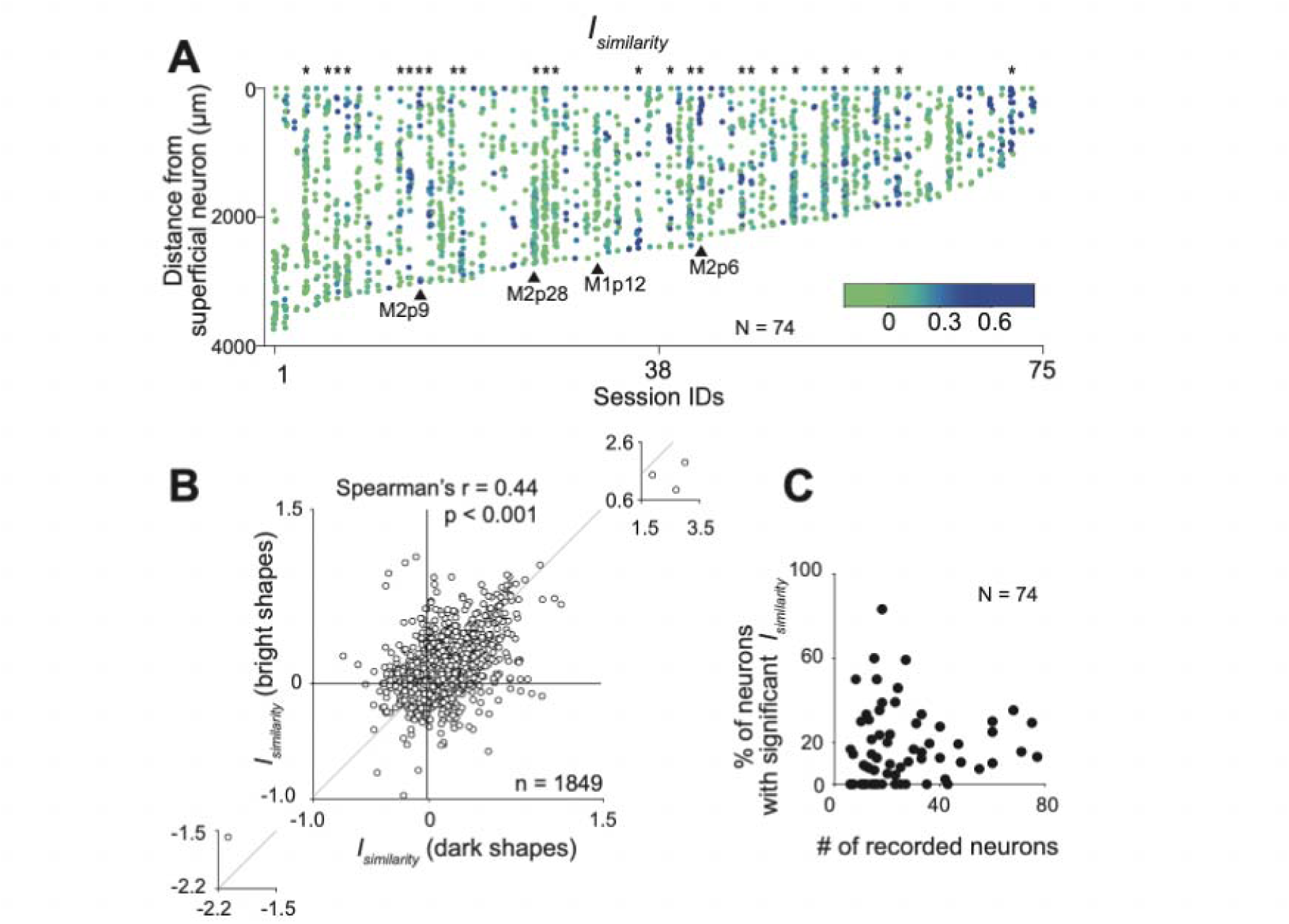
Population results for clustered shape tuning. **(A)** *I_similarity_* (color) for each neuron is illustrated as a function of recorded depth (Y axis) across sessions (columns) sorted according to the probe length over which neurons were recorded. To ensure clarity, green to blue span the 0 to 0.6 range of *I_similarity_*; values outside of this range were set to either green (for *I_similarity_* < 0) or blue (for *I_similarity_* > 0.6). Asterisks above the panels indicate penetrations with 5 or more neurons with significant *I_similarity_*. Penetrations depicted in Figures 2, 3, 4 and 8 are identified. **(B)** Intercept of regression lines (*I_similarity_*) based on tuning similarity for dark (X axis) versus bright shape stimuli (Y axis). Results from the two sets of stimuli are consistent. **(C)** Percentage of neurons with *I_similarity_* significantly different from 0 (Y axis) as a function of the number of simultaneously studied neurons (X axis). N denotes number of penetrations.

Neurons with high similarity in tuning (Figure 5A) were widely distributed across 52 of the 74 penetrations in small, sparse clusters (blue). Only 26 penetrations had 5 or more neurons with significant *I_similarity_* (Figure 5A asterisks). When the number of recorded neurons was > 30, the percentage of neurons with significant *I_similarity_* asymptoted to ∼20% (Figure 5C) suggesting that shape clusters were not “columnar” extending across laminae in V4, and simultaneous recordings from > 30 neurons (achieved on 20/74 penetrations) were required to provide a fair assessment of the fraction of neurons that were similarly shape-tuned. Results were consistent between the two animals (not shown).

### Control experiments

One reason for the sparsity of similarly tuned clusters may be our stimulus protocol. Different V4 neurons within a penetration can have overlapping but distinct RF profiles, and because our shape silhouettes were presented in one position (at the center of the aggregate RF), they may not have provided identical stimulation (in terms of RF overlap) for all neurons, and thus may not be well suited to assess similarity in tuning of simultaneously recorded neurons. To consider this possibility rigorously, we used a set of spatially homogeneous textures to assess tuning similarity (Figure 6). We also assessed how shape tuning similarity changed with position for a subset of our penetrations (Figure 8 and 9).

**Figure 6.**
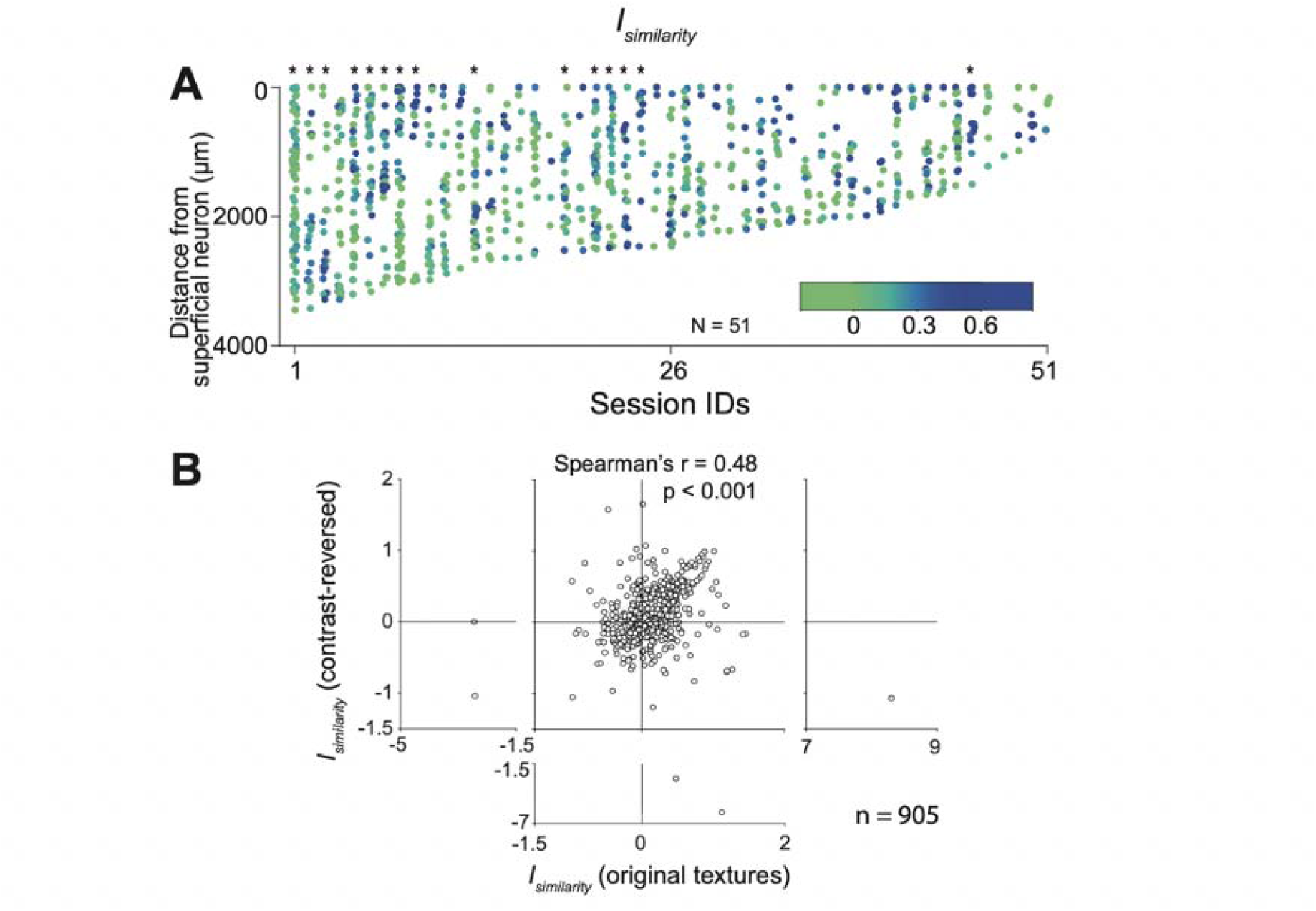
Population results for clustered texture tuning. **(A)** *I_similarity_* of texture tuning similarity for each neuron (color) is illustrated as a function of recorded depth (Y axis) across 51 penetrations (columns). Blue and green dots identify neurons with and without high texture tuning similarity with neighboring neurons respectively (see details as in Figure 5A). Asterisks above the panel indicate penetrations with 5 or more neurons with significant *I_similarity_*. **(B)** *I_similarity_* from regression lines for tuning similarity based on responses to original textures (X axis) versus contrast-reversed textures (Y axis). Results from the two sets of stimuli are consistent.

### Population results: texture tuning

We studied the responses of 979 neurons across 51 sessions (M1:17; M2:34) using a set of 40 texture stimuli and their contrast-reversed counterparts (Figure 1C). Texture stimuli were homogeneous in terms of their image statistics and sized to overlap the RFs of all manually mapped channels (see Materials and Methods). This ensured that the RF of every neuron was stimulated with the same image statistics, avoiding the shortcoming of shape stimuli which may have overlapped a given neuron’s RF profile differently than another neuron’s RF. We assessed similarity in tuning across all pairs of neurons recorded within a penetration (ranging from 6 to 83 per penetration). As with shape stimuli, we found that a small subset of neurons (204/979; M1:18; M2:186) across 35/51 (M1:8; M2:27) penetrations showed clustered selectivity, i.e. evidence of declining tuning similarity with increasing inter-neuron distance. Figure 6A shows the *I_similarity_* of regression lines for texture tuning similarity (color) as a function of recorded depth (Y axis) across 51 penetrations (columns). Across the 51 penetrations, 15 penetrations had 5 or more neurons that exhibited regression lines with statistically significant intercepts (asterisks, Figure 6A). Our results from texture (black symbols in Figure 7) are consistent with those from shape (gray symbols in Figure 7): the fraction of neurons with a significant intercept asymptote to ∼20% when the number of recorded neurons was large. Tuning similarity based on both sets of texture data (original and contrast-reversed) were highly consistent (Figure 6B). These results confirm that the sparse occurrence of tuning similarity observed above with shape stimuli cannot be dismissed as resulting from differential stimulation of the RFs of neurons by our shape stimuli.

**Figure 7.**
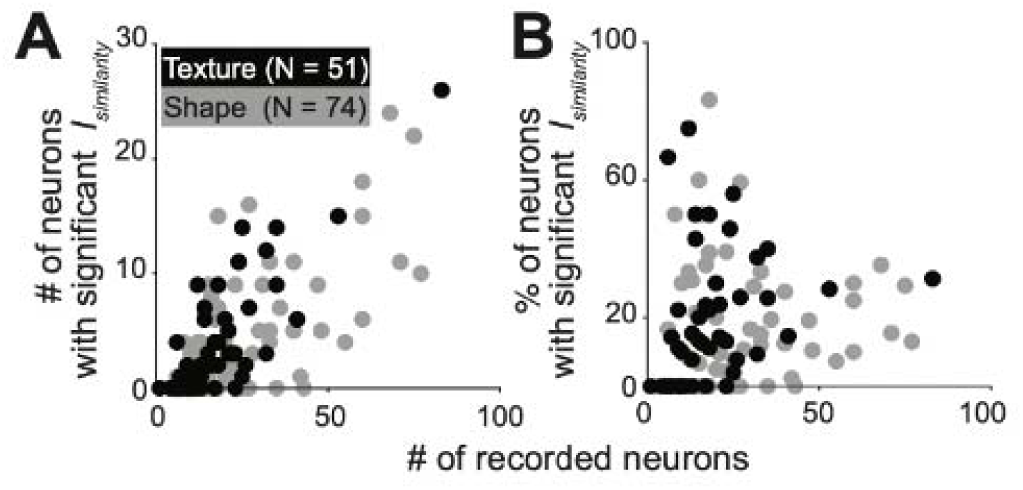
Effect of recording yield size on the detection of clustered tuning. Number (A) and percentage (B) of neurons with *I_similarity_* significantly different from 0 (Y axis) as a function of the number of simultaneously studied neurons (X axis). N denotes number of penetrations. Black and gray circles indicate penetrations where responses to shape and texture responses were studied.

### Position invariance for shape tuning similarity

To further evaluate how differential stimulation of neuronal RFs (due to the variability of RF profiles across neurons, see Figures 8B and 8D) influenced our shape tuning similarity assessments, in 34 sessions we studied responses to a subset of 15 stimuli at 5 positions – at the aggregate RF center and at four positions displaced ½ × RF diameter (north, south, east and west) relative to the center (Figure 8B). We then asked whether tuning similarity profiles were consistent across positions. Results from two representative recording sessions are shown in Figures 8A and C, and in both cases, tuning similarity patterns were similar across all five positions: a small cluster of similarly tuned neurons is evident across all positions in Figure 8A, and no such cluster emerged at any position in Figure 8C. Next, we evaluated whether tuning similarity across nearby neurons was greater when comparing responses at the optimal RF position for each neuron. For each neuron, we first identified the RF location at which our shape stimuli evoked the most diverse responses (see RF localization in Materials and Methods) and then assessed tuning similarity based on responses at these optimal locations (Figure 9). Tuning similarity patterns on the basis of the most diverse responses were similar to that quantified based on responses at the aggregate RF center (compare Figure 9A left vs 8A, and Figure 9A right vs 8C). The similarity matrices based on reduced stimulus sets were also similar to that from the main experiment with 120 stimuli (not shown). We found similar results across the 34 penetrations (n = 723) where we conducted the position tests. To summarize this result across penetrations, we assessed tuning similarity based on responses to stimuli at the aggregate RF center and at optimal RF position for each penetration (as in Figure 9A). We then constructed regression lines and quantified *I_similarity_* for each neuron for the RF center vs RF optimal conditions. We found a statistically significant positive correlation between *I_similarity_* in the RF aggregate center versus RF optimal positions (r = 0.349, p < 0.0001, Figure 9B). These results provide evidence against the possibility that stimulus position and RF variability contributed to the sparsity of clusters observed with our shape stimuli.

**Figure 8.**
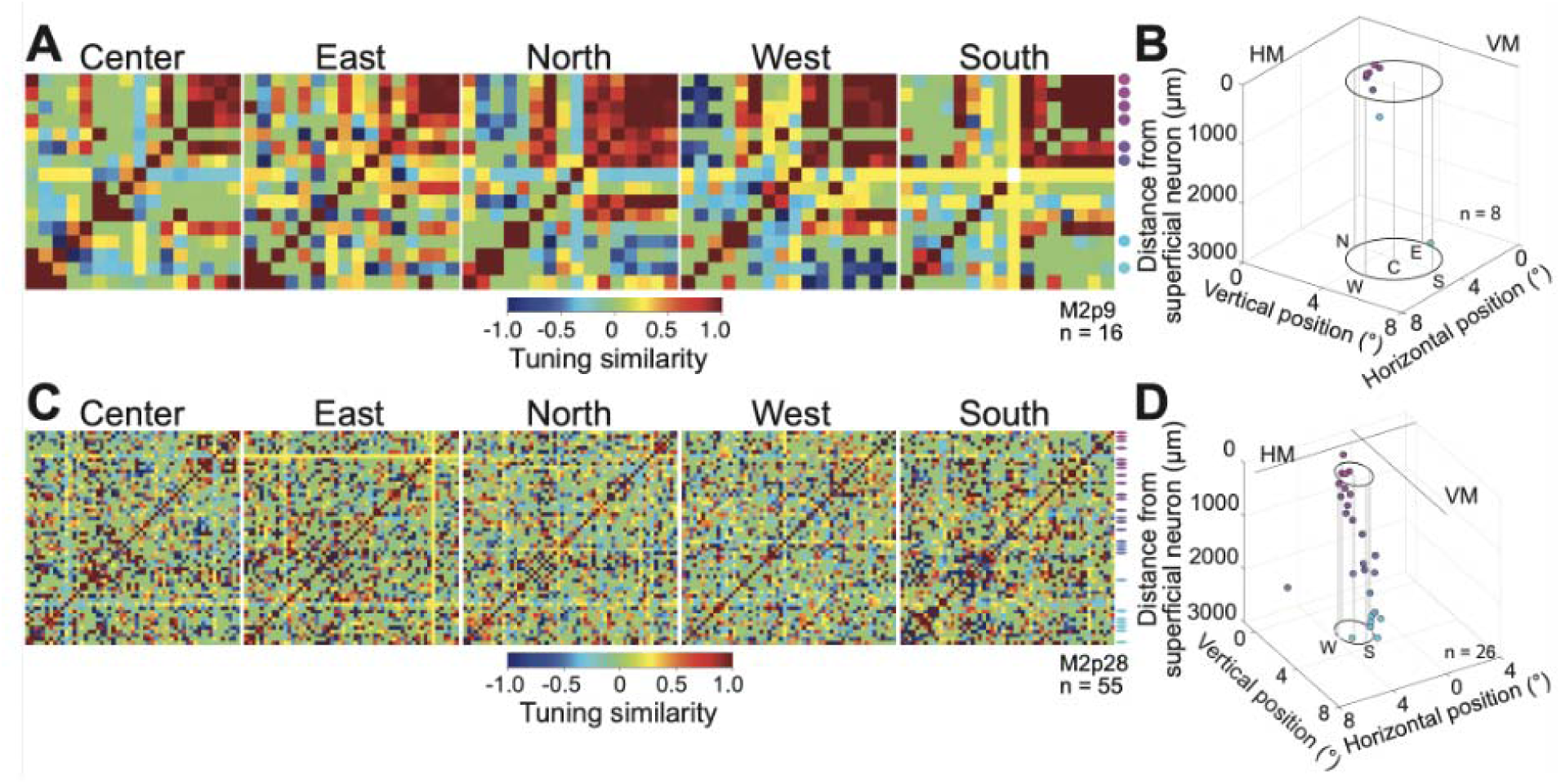
Position invariance of tuning similarity. **(A)** Example penetration with a small cluster of similarly tuned neurons (M2p9). Similarity matrices based on the responses to 15 shape stimuli presented at five positions relative to aggregate RF center are shown. Colored dots at right identify neurons that showed visually evoked responses during preliminary characterization (see B). **(B)** RF center position (X and Y axes) for the neurons in this recording session plotted as a function of the depth from superficial neuron (Z axis). RF centers were estimated from automated RF localization procedure (see Materials and Methods). Eight neurons that failed to yield an RF estimate are not depicted. Black circles: aggregate RF. The five stimulus positions (Center, North, East, West and South) are indicated by vertical gray lines. HM: Horizontal meridian. VM: Vertical meridian. Inset n indicates neuron numbers with quantified RF centers. **(C)** Similarity matrices from a penetration (M2p28) without a cluster of similarly tuned neurons. All other details as in A. **(D)** RF position (X and Y axes) and depth (Z axis) of neurons recorded for the penetration depicted in C. Details as in B.

**Figure 9.**
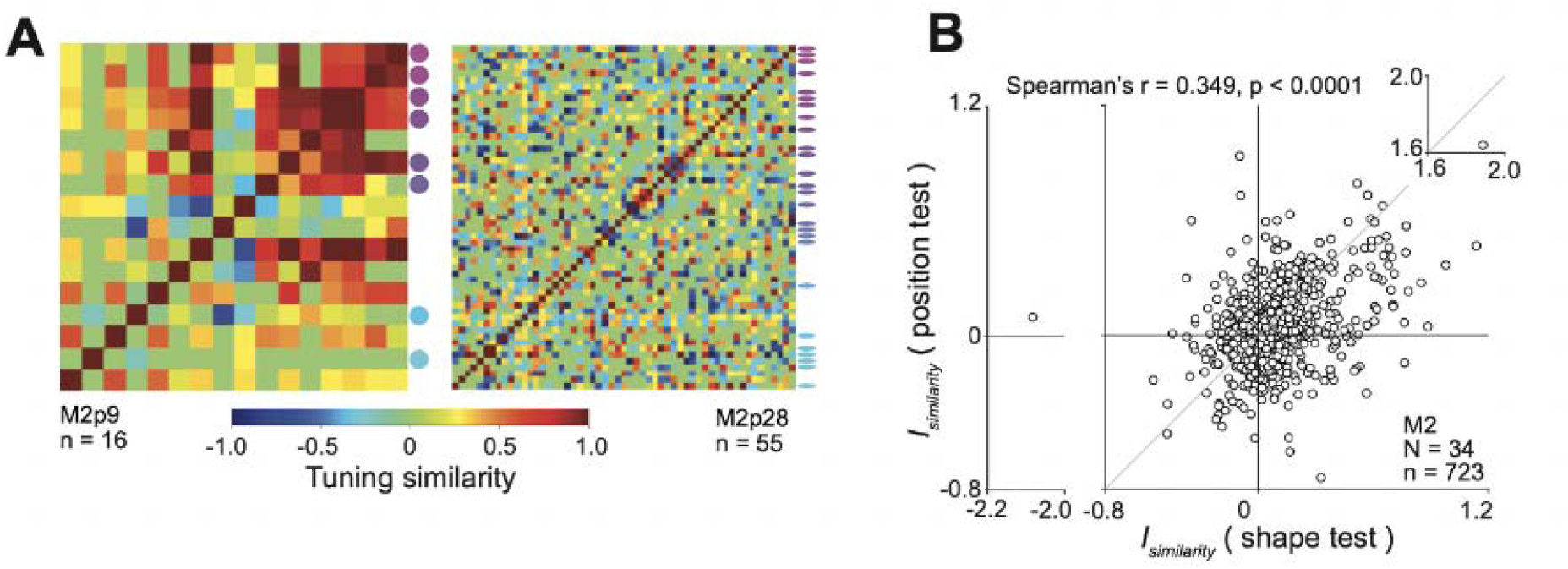
Tuning similarity based on responses at best RF position. **(A)** Single neuron optimized measure of tuning similarity for the two penetrations depicted in Figures 8AB (left) and 8CD (right). For each neuron in each penetration, we first identified the optimal stimulus position as the one associated with the greatest dispersion in responses. We then used the responses from the optimal position to construct a tuning similarity matrix (see Materials and Methods). **(B)** *I_similarity_* computed in the position test (Y axis) vs *I_similarity_* computed in the main shape test (X axis). Inset N and n indicate penetration and neuron numbers, respectively.

### Simulation of tuning progression

Across our data set, we often recorded neurons over a probe length > 2000 μm, suggesting that many of our penetrations were oblique rather than radial. If V4 did include columns for shape tuning, then an oblique penetration may mean gradual (or abrupt) changes in tuning preference along the length of the probe. We conducted simulations to visualize the tuning similarity matrices we might observe for different patterns of tuning progression across the length of the probe for oblique penetrations. We used the angular position × curvature model (APC model from Pasupathy and Connor, 2001) to simulate the responses of 30 model V4 neurons to the 120 shape stimuli we used, and used them to visualize similarity matrices resulting from a non-radial trajectory of our probe through columns of neurons with similar tuning. We considered the possibilities that nearby neurons along the length of our probe exhibited a gradual change in tuning for curvature alone (Figure 10A), angular position alone (Figure 10B) or both (Figure 10C). The resulting similarity matrices (Figures 10A, B, and C, bottom panels) were quite unlike our V4 data (compare with Figures 3 and 4). Random choice of tuning peaks in the APC space produced similarity matrices reminiscent of the majority of V4 data (compare Figure 10D bottom with Figure 4). When we interleaved the random choice of tuning peaks with short stretches of similar tuning (Figure 10E), the resulting similarity matrices matched our observation of localized clusters (compare Figure 10E bottom with Figure 3). These results support the idea that V4 does not exhibit columnar tuning for shapes. Rather, the observed patterns correspond best to random selectivity interspersed with small domains of similar tuning for shapes.

**Figure 10.**
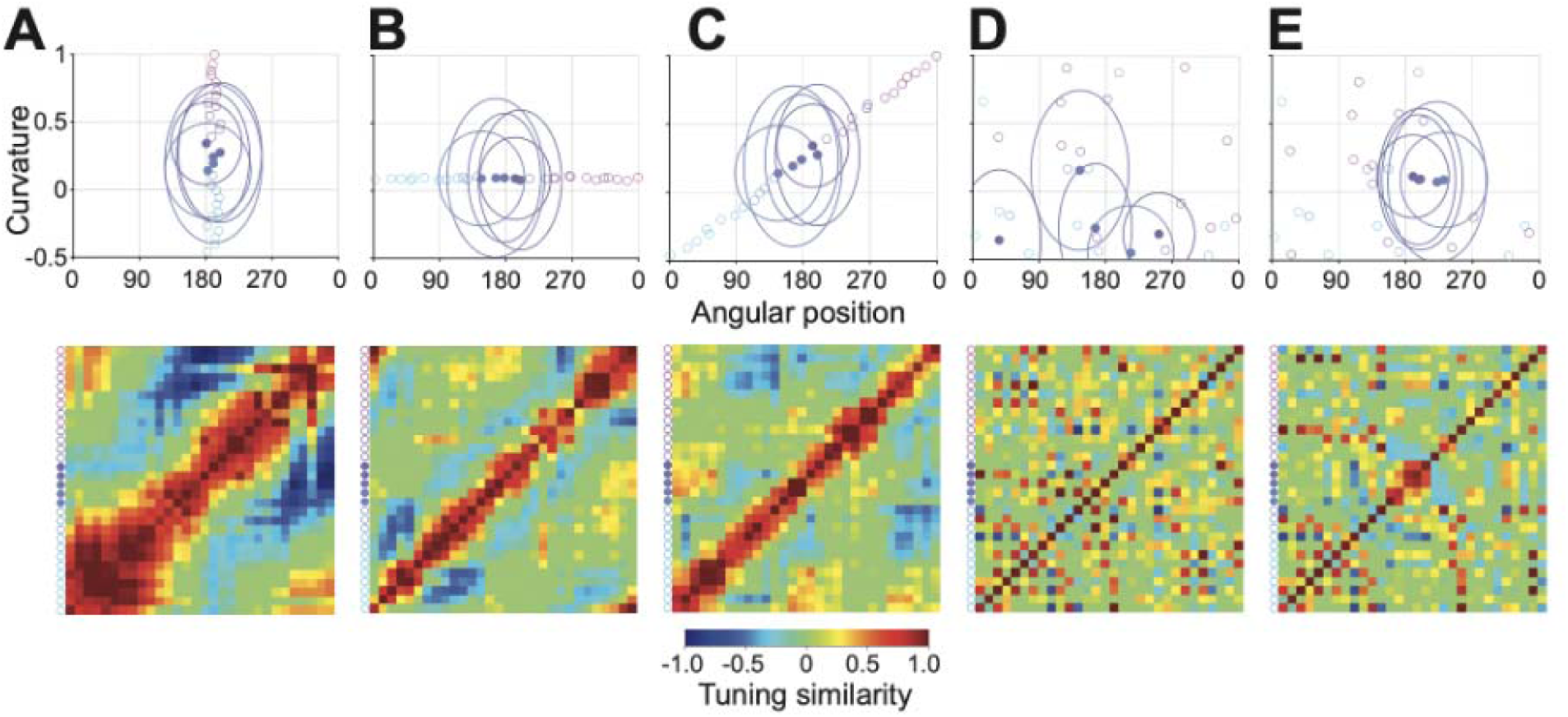
Simulation: tuning progression and the resulting tuning similarity patterns. **(A-E)** The top row shows shape tuning peaks of model V4neurons in the angular position × curvature space. Responses of model units to shape stimuli were dictated by a 2D Gaussian function centered at the peak position shown. Curvature runs from -0.5 (moderate concavity) to 1.0 (sharp convexity) and specifies the curvature of the preferred boundary feature. Angular position runs from 0° (pointing right) in a counterclockwise direction and specifies the position of the preferred feature relative to object center. Different tuning peak progressions (left to right) across 30 model neurons along the length of the probe (purple: superficial; cyan: deep) were considered for each panel. Tuning similarity matrices across model neurons are shown at the bottom. Tuning similarity ranges from -1 (blue: opposite preference) to 1 (red: identical preference). The tuning similarity matrices were constructed by computing noise-corrected correlation coefficient between modeled spike counts of 30 neurons to 120 shapes used in the main experiment. For each of 120 shapes, noisy spike counts were modeled for 3 trials repetitions by implementing Poisson spiking (see Materials and Methods). **(A)** Tuning preference along the curvature dimension (Y axis) gradually shifts from a preference for sharp convexity (curvature = 1) to medium concavity (-0.5) from superficial to deep neurons, while the preferred angular position remained at 191° (pointing left). All other parameters of the 2D Gaussian function (SD and amplitude, denoted by circles) were randomly set across neurons. **(B)** Tuning preference along the curvature (Y axis) remains constant at low convexity with small fluctuation along the curvature, while the preferred angular position gradually shifts from a preference for features pointing to the right in a counterclockwise direction. All other details as in A. **(C)** Tuning preference of modeled neurons gradually changes along both the angular position and curvature dimensions. Step sizes of tuning shifts for angular position and curvature were the same as those in A and B. **(D)** Preferred angular position and curvature were chosen at random for individual model neurons. **(E)** Five neurons exhibiting systematic shift of preferred angular curvature from left (180°) in a counterclockwise direction were interleaved in neurons with random peaks.

## Discussion

We used high-density Neuropixels probes to investigate whether V4 neurons across laminae share similar tuning for stimulus features. We found that ∼20% of neurons share shape and texture tuning with nearby neurons, but such clusters seldom encompass all laminae. This was the case despite the fact that neurons were driven by shape stimuli. Our results cannot be explained on the basis of stimulus choice, diversity in RF stimulation or oblique trajectories. We conclude that V4 functional organization for shape and texture may only be at a categorical level and the lack of columnar structure may reflect the variety of inputs integrated by V4 neurons to build high-dimensional properties (Purves et al.,1992). Finally, lack of large clusters of similarly tuned neurons may explain the lack of a strong effect of V4 microstimulation on perception (Dagnino et al., 2015).

### Relationship to prior studies

Many studies have investigated functional organization in V4 with imaging methods. In a recent study, Srinath and colleagues demonstrated segregated modules encoding shape stimuli defined by 2D versus 3D cues (Srinath et al., 2021). Intrinsic signal imaging has also revealed functional domains specialized for processing curvature versus rectilinear stimulus features (Hu et al., 2020), motion direction (Li et al., 2013) and high versus low spatial frequencies (Lu et al., 2018). In all of these studies, the focus has been categorical organization – 2D versus 3D cues, curved versus rectilinear shapes, high versus low spatial frequency, etc. – rather than fine-scale organization, i.e. investigations into whether the profile of the tuning curve along a specific feature dimension is similar for nearby neurons. Furthermore, because these studies were based on imaging, they are biased toward characterizing the upper layers.

When studies could examine tuning similarity, e.g. with multiphoton imaging of single neuron responses, or with electrode recording across layers, authors have found considerable diversity in the tuning of nearby neurons (Jiang et al., 2021; Tang et al., 2020). For example, using electrode recordings, Li and colleagues found consistency in categorical organization across layers, i.e. whether neurons are direction selective or not, but considerable diversity in tuning preference of individual neurons within domains (2013). These results are largely consistent with studies investigating functional architecture for color in V4, where authors have investigated fine-scale similarity in tuning with electrode recordings spanning all layers (Schein et al., 1982; Yoshioka and Dow, 1996; Zeki, 1973). While results across studies are mixed, based on the region of focus and the definition of color, a comprehensive evaluation by Kotake et al.(2009) found sporadic clusters and weak evidence for columnar organization.

We investigated fine-scale tuning similarity for shape and texture across laminae, analogous to electrode recording investigations and orthogonal to much of the prior imaging work. Our results are consistent with prior work discussed above, in that we also find considerable diversity in the fine-scale tuning of nearby neurons. We do find small clusters of similarly tuned neurons–roughly 20% of neurons occupy such clusters– but these clusters are spatially confined and do not extend across all layers. Given that we used simple 2D silhouettes and texture patches, one may wonder if more complex/naturalistic shape stimuli could reveal columnar structure. We do not think so. Higher-dimensional stimuli are likely to be encoded more sparsely given the high-dimensional tuning of V4 neurons. On the other hand, low-dimensional stimuli such as oriented gratings reveal columns but only close to foveal representation in V4 (Ghose and Ts’o, 1997).

### Origin of columns or the lack thereof

A lack of columns for shape and texture tuning in V4 may seem surprising given the observation of detailed functional columnar structure in V1 and MT, and the demonstration that all of cortex is organized as a set of ontogenetic columns (Rakic, 1988) that run from white matter to the pial surface. However, across species and brain areas, many instances of the absence of any clustering (based on physiological properties) or local clustering without columnar structure have been previously documented (Horton and Adams, 2005). For example, mice and rats lack orientation columns in the primary visual cortex and the organization is “salt and pepper” (Ohki et al., 2005). This is also true of the squirrel, an animal with high visual acuity and many orientation-tuned neurons (Van Hooser et al., 2005). Another striking example is the organization in auditory cortex in cat and rodent, where overlapping maps of multiple stimulus features have been documented in thalamo-recipient layers III and IV (Linden and Schreiner, 2003), yet there is also considerable heterogeneity in best frequency responses across layers (Tischbirek et al., 2019).

Current evidence suggests that emergence of columns in primary visual cortex depends not on the visual acuity of the animal in question, or the size of V1, but rather on the retino-cortical mapping ratio (Jang et al., 2020). For example, the ferret, tree shrew, gray squirrel and rabbit all have a similar sized V1, but columns are evident only in the former two animals with a smaller retino-cortical ratio. In other words, when the cortical representation is associated with a large expansion, columns emerge even when wiring is established by statistical sampling (Ringach, 2004, 2007). Then, by analogy, one could speculate that the lack of columnar structure with our stimuli in V4 may relate to the contraction in the V1/V2 to V4 representation (due to the smaller size of V4). By this same analogy, we would expect larger clusters in V2 (given the larger size) and our preliminary results support this hypothesis (Kim et al., unpublished observations). Furthermore, V4 receives a great diversity of inputs encoding form, color and motion direction, and individual V4 neurons exhibit nonlinear tuning in a high-dimensional space (Pasupathy et al., 2019, 2020). Purves and colleagues have hypothesized that such complex integration of signals from multiple pathways dilutes the coherence in activity of neighboring neurons, making column formation less likely (Purves et al., 1992). Furthermore, if different sources of input target different V4 layers, columnar structure would be less likely. While it seems paradoxical that columnar structure is evident in MT given its small size, this may be because MT receives far fewer inputs than V4 (Felleman and Van Essen, 1991) and these inputs are less diverse, such that MT tuning remains low-dimensional. We concede that these arguments are speculative and further experiments are needed to investigate the raison d’etre of columnar structure.

## Conclusion

Overall, based on our work we conclude that functional organization is not columnar for shape and texture stimuli in V4, reflecting the number and diversity of inputs integrated to build the different types of selectivity, e.g. for color, shape, texture. Prior studies suggest that there may be organization at the categorical level and if this is indeed true, then future V4 perturbation studies could be most impactful if they target categorical rather than fine scale discriminations.

## Acknowledgements

This work was supported by NEI grant R01 EY018839 and R01 EY029601 to A.P., Vision Core Grant P30EY01730 to the University of Washington, Office of Research Infrastructure Programs Grant OD010425 to the Washington National Primate Research Center and JSPS KAKENHI Grant 23K06785 to TN, JST ERATO JPMJER1801, and CREST JPMJCR18A5. We thank Taekjun Kim for help with visual stimuli, Amber Fyall for animal care and behavioral training, WaNPRC Instrumentation Services for hardware support, and Dina Popovkina, Zhiwen Ye, Nick Steinmetz, Greg Horwitz, Taekjun Kim, Dean Pospisil and Rohit Kamath for helpful discussions and comments on the manuscript.

